# Revisiting the Global Workspace: Orchestration of the functional hierarchical organisation of the human brain

**DOI:** 10.1101/859579

**Authors:** Gustavo Deco, Diego Vidaurre, Morten L. Kringelbach

**Affiliations:** Center for Brain and Cognition, Computational Neuroscience Group, Department of Information and Communication Technologies, Universitat Pompeu Fabra, Roc Boronat 138, Barcelona, 08018, Spain; Institució Catalana de la Recerca i Estudis Avançats (ICREA), Passeig Lluís Companys 23, Barcelona, 08010, Spain; Department of Neuropsychology, Max Planck Institute for Human Cognitive and Brain Sciences, 04103 Leipzig, Germany; School of Psychological Sciences, Monash University, Melbourne, Clayton VIC 3800, Australia; Department of Psychiatry, University of Oxford, Oxford, UK; Center for Music in the Brain, Department of Clinical Medicine, Aarhus University, DK

## Abstract

A central, unsolved challenge in neuroscience is how the brain orchestrates function by organising the flow of information necessary for the underlying computation. It has been argued that this whole-brain orchestration is carried out by a core subset of integrative brain regions, commonly referred to as the ‘global workspace’, although quantifying the constitutive brain regions has proven elusive. We developed a *normalised directed transfer entropy* (NDTE) framework for determining the pairwise bidirectional causal flow between brain regions and applied it to multimodal whole-brain neuroimaging from over 1000 healthy participants. We established the full brain hierarchy and common regions in a ‘functional rich club’ (FRIC) coordinating the functional hierarchical organisation during rest and task. FRIC contains the core set of regions, which similar to a ‘club’ of functional hubs are characterized by a tendency to be more densely functionally connected among themselves than to the rest of brain regions from where they integrate information. The invariant *global workspace* is the intersection of FRICs across rest and seven tasks, and was found to consist of the precuneus, posterior and isthmus cingulate cortices, nucleus accumbens, putamen, hippocampus and amygdala that orchestrate the functional hierarchical organisation based on information from perceptual, long-term memory, evaluative and attentional systems. We confirmed the causal significance and robustness of this invariant global workspace by systematically lesioning a generative whole-brain model accurately simulating the functional hierarchy defined by NDTE. Overall, this is a major step forward in understanding the complex choreography of information flow within the functional hierarchical organisation of the human brain.

## Introduction

Over the last decades, careful research within systems neuroscience has suggested that the brain is hierarchically organised in terms of anatomical structure (Bullmore and Sporns, 2012; Felleman and Van Essen, 1991; Hagmann *et al.*, 2008; Markov *et al.*, 2014; Mesulam, 1998; van den Heuvel and Sporns, 2011; Zamora-Lopez *et al.*, 2010) and function (Atasoy *et al.*, 2016; Atasoy *et al.*, 2017; Buckner and DiNicola, 2019; Huntenburg *et al.*, 2018; Margulies *et al.*, 2016). Nevertheless, a key remaining challenge is to determine how precisely this hierarchical organisation allows the brain to orchestrate function by organising the flow of information and the underlying computations necessary for survival.

A large body of research has argued that whole brain orchestration is likely to be carried out by a core subset of integrative brain regions. For example, according to the classic model of Norman and Shallice (Norman and Shallice, 1980), this processing involves the prefrontal cortices in charge of the supervisory attentional regulation of lower-level sensori-motor chains. In contrast, Baars proposed the concept of a ‘global workspace’, where information is integrated in a small group of brain regions before being broadcasts to many other regions across the whole brain (Baars, 1989). Extending this framework, Dehaene and Changeux (1998) proposed their ‘global neuronal workspace’ hypothesis that associative perceptual, motor, attention, memory, and value areas interconnect to form a higher-level unified space where information is broadly shared and broadcast back to lower-level processors. Colloquially, the global workspace is thus akin to a small core assembly of people in charge of an organisation. Larger brain network organisation has been shown to be efficient, robust and largely fault tolerant (Alstott *et al.*, 2009; Bullmore and Sporns, 2012; Honey and Sporns, 2008), yet the effects of lesioning such a core assembly are currently unknown.

Until now, a key obstacle to advancing our understanding of the human brain’s functional hierarchical organisation has been the lack of suitable whole-brain measurements. However, the advent of ‘big data’, such as Human Connectome Project (HCP) (Glasser *et al.*, 2016b; Van Essen *et al.*, 2013), has created large multimodal whole-brain neuroimaging datasets of healthy individuals both in resting state and whilst performing many different tasks. Potentially, the development of more advanced neuroimaging methods could allow for the estimation of bi-directional flow of information between all regions across the whole brain, which could subsequently be used to characterise the functional hierarchical organisation of the brain.

If the precise characterisation of hierarchical information flow could be obtained, this would allow for the discovery of a core set of brain regions responsible for integration and orchestration. We propose here the ‘Functional Rich Club’ (FRIC) as the core set of regions, separate from the rest of the brain, which similar to a ‘club’ of functional hubs are characterized by a tendency to be more densely functionally connected among themselves than to other brain regions from where they receive integrative information. This notion is related to previous static descriptions of the *anatomical* rich club (van den Heuvel and Sporns, 2011; Zamora-Lopez *et al.*, 2010), which includes nodes in a network with a tendency for high-degree nodes to be more densely connected among themselves than nodes of a lower degree. However, FRIC is a dynamic measure based on bidirectional flow of information that is not constrained by anatomy and thus will change across different tasks. Using our colloquial example of a core assembly, for different tasks some people remain through all executive meetings while others are substituted in and out based on their expertise. In a similar manner, FRIC would include both common and task-specific brain regions as a result of the different flow of information for different kinds of tasks. Following the original ideas of Baars, we propose that the invariant ‘global workspace’ is the intersection of the different sets of task-related FRICs.

Here, we apply our novel *normalised directed transfer entropy* (NDTE) framework on state-of-the-art data from over 1000 healthy HCP participants to discover the global workspace given as the intersection of FRICs from seven different tasks and resting condition. The NDTE framework provides a bidirectional causal description of the functional information flow underlying brain signals using a normalised version of the transfer entropy with appropriate surrogate methods and aggregation of *p*-value statistics across the many participants (see Methods).

Furthermore, we validate the causal significance of the invariant global workspace through constructing and selectively lesioning a whole-brain model that accurately simulates the empirical functional hierarchy and thus describes the underlying dynamical mechanisms. Systematic lesioning of subsets of regions in the FRIC in this model establishes the causal role of the global workspace in orchestrating function and allows us to characterise the efficiency and robustness. Overall, this is a major step forward in understanding the complex choreography of information flow within the functional hierarchical organisation of the human brain.

## Results

The overall aim is to find the regions orchestrating the functional hierarchical organisation, sometimes called the ‘global workspace’ (Baars, 1989; Dehaene *et al.*, 1998). In order to obtain robust results, we used multimodal neuroimaging data from 1003 normal participants (Human Connectome Project, HCP) whilst performing seven tasks and resting state (Glasser *et al.*, 2016b). As sketched in **Figure 1**, this is made possible by the framework of Normalised Directed Transfer Entropy (NDTE), which allows for the precise characterisation of the causal flow of information between pairs of brain regions and consequently the estimation of the overall hierarchical organisation of the human brain in a given state. Furthermore, this quantitative framework can be used to identify the smallest core set of brain regions that integrate and orchestrate function in a given task. This ‘Functional Rich Club’ (FRIC) comprises the core set of regions, which are separate from the rest of the brain, and which is defined as a ‘club’ of functional hubs that are characterized by a tendency to be more densely functionally connected among themselves than to other brain regions from where they receive integrative information.

**Figure 1.**
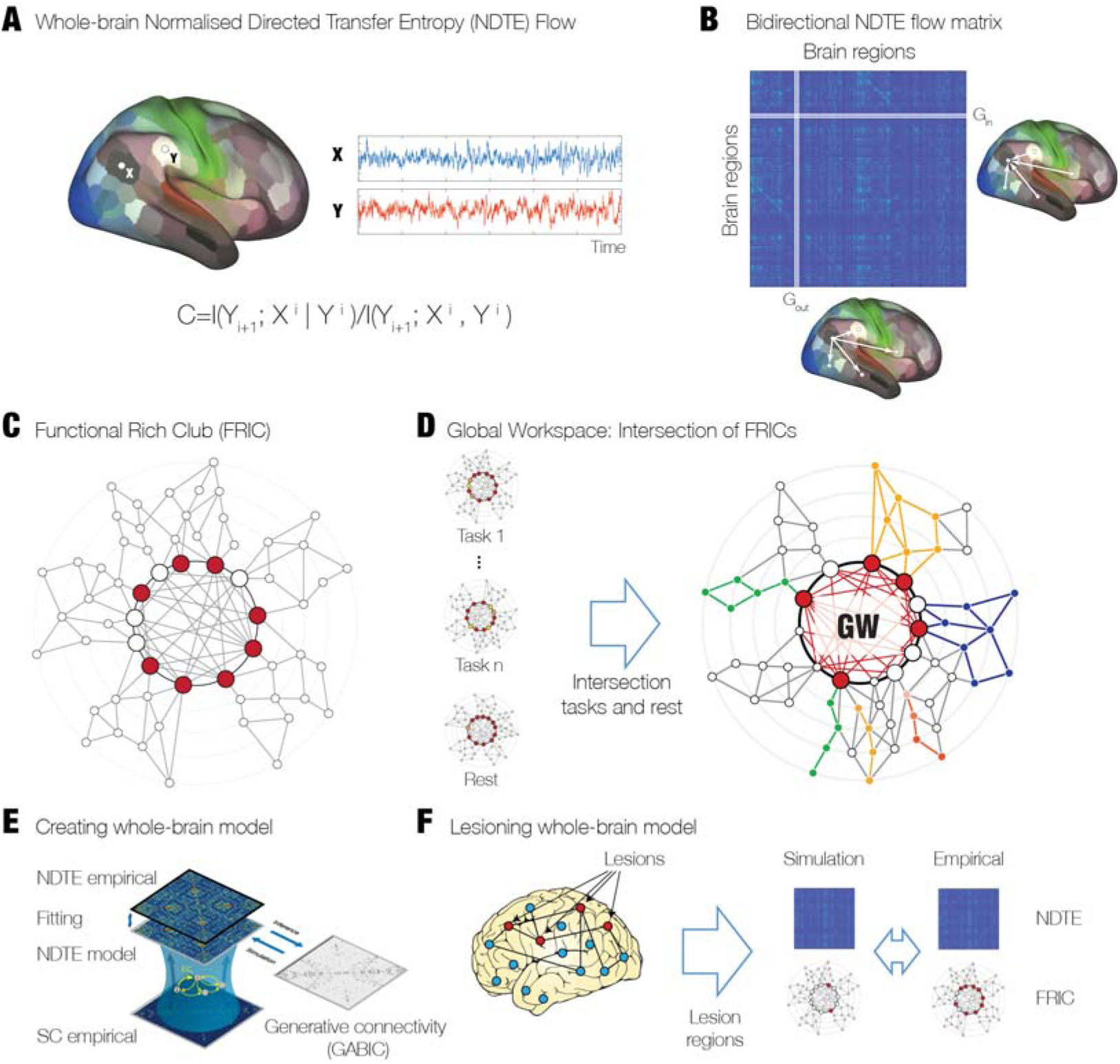
Overview of general theoretical framework. **A)** Causal bidirectional flow of information between any two brain regions is determined by computing the pairwise normalised directed transfer entropy (NDTE). The statistical significance is determined at the individual level by using the circular time shifted surrogates method (Quiroga et al., 2002) and at the group level by using P-level aggregation across individuals. **B)** The functional hierarchical organisation is given by the regions. For each brain region, the total incoming flow of information, G_in_, is simply the sum of all full NDTE matrix, where the rows contain the target regions and the columns contain the source sources (ie the sum over the rows in the matrix). Similarly the total outgoing flow of information, G_out_, is the sum over all targets (ie the sum over the columns). **C)** The functional rich club (FRIC) is the smallest set of brain regions that integrate and orchestrate function in a given task. It can be identified as the most highly connected brain regions that 1) are more densely connected within themselves than to regions with lower connectivity, whilst 2) having the highest level of incoming directed flow (G_in_) and 3) the lowest outgoing directed flow (G_out_, see Methods). **D)** The global neuronal workspace has to be relevant to all tasks and situation and must therefore be the common FRIC members across many different tasks, ie the intersection of FRICs from tasks and rest. **E)** In order to establish the causal importance of the FRIC, we fit a whole-brain model to the resting NDTE empirical data and extract the underlying effective connectivity (see Results and Methods). **F)** The whole-brain model is then systematically lesioned for regions belonging to the FRIC and compared to lesioning non-FRIC members. Overall, this confirms the causal importance of these regions in the orchestration of the functional hierarchical organisation of the human brain.

Finally, we identify the ‘global workspace’ as the intersection of the FRIC members across tasks and rest. The key functional relevance of these core set of brain regions is then shown to be causal through creating and lesioning a mechanistic whole-brain model that can generate the NDTE flow during resting. Lesioning the FRIC members in this whole-brain model causes significant breakdown of information flow compared to lesioning non-FRIC members, attesting to the major significance of the global workspace in orchestrating functional organisation.

### Functional hierarchical organisation of resting

We characterised the functional hierarchical organisation of the resting state of 1003 HCP participants. For this purpose, we extracted the causal bidirectional flow of information between brain regions using the concept of normalised directed transfer entropy (NDTE) (see Methods), which is an information theoretical measure of causality between two time series. This allows us to infer the underlying bidirectional causal reciprocal communication between any source and target regions. Specifically, this computes the mutual information directed flow, i.e. the predictability of a target in the future given the past of the source region, beyond the predictability from its own past (see Equation 1 in Methods). This is then normalised by the mutual information that both source and target have about the future of the target (see Equation 8 in Methods). At the individual level, we compute the statistical significance by using the circular time shifted surrogates method which has been shown to be particularly well-adapted to causal measurements (Quiroga et al., 2002). At the group level, we aggregate the p-values corresponding to each pairwise NDTE flow by using the Stouffer method (Stouffer *et al.*, 1949) (see Methods).

We computed two matrices containing the causal NDTE flow between the regions in two different parcellations (see Methods). For a fine-scale parcellation, we used a modified version of the Glasser parcellation with a total of 378 regions (360 cortical and 18 subcortical regions) (Glasser *et al.*, 2016a). For a coarser-scale parcellation suitable for whole-brain computational modelling, we used a modified Desikan-Killiany parcellation which included subcortical regions (DK80, 62 cortical regions and 18 subcortical regions) (Desikan *et al.*, 2006; Klein and Tourville, 2012).

In order to establish the hierarchical organisation, we compute the total incoming and outgoing information for all brain regions. More specifically, for each brain region, the total incoming flow of information, G_in_, is the sum of all sources (ie the sum over the rows in the matrix). Similarly, the total outgoing flow of information G_out_, is the sum over all targets (ie the sum over the columns). We also compute the total information being processed in a brain region as the sum: G_tot_= G_in_ + G_out_, which is possible given that the measures have been properly normalised (see above and Methods).

We show the functional hierarchy described by each of the incoming (G_in_), outgoing (G_out_) and total (G_tot_) directional information flow computed from the NDTE matrix of 1003 HCP participants (see **Figure 1** and Methods) in the Glasser parcellation (**Figure 2A**) and DK80 (**Figure 2D**). As can be seen (especially in the enlarged version in **Figure S1**), the outgoing information flow, G_out_, is highest in sensory areas, while incoming information, G_in_, is highest in higher-order, integrative transmodal areas.

**Figure 2.**
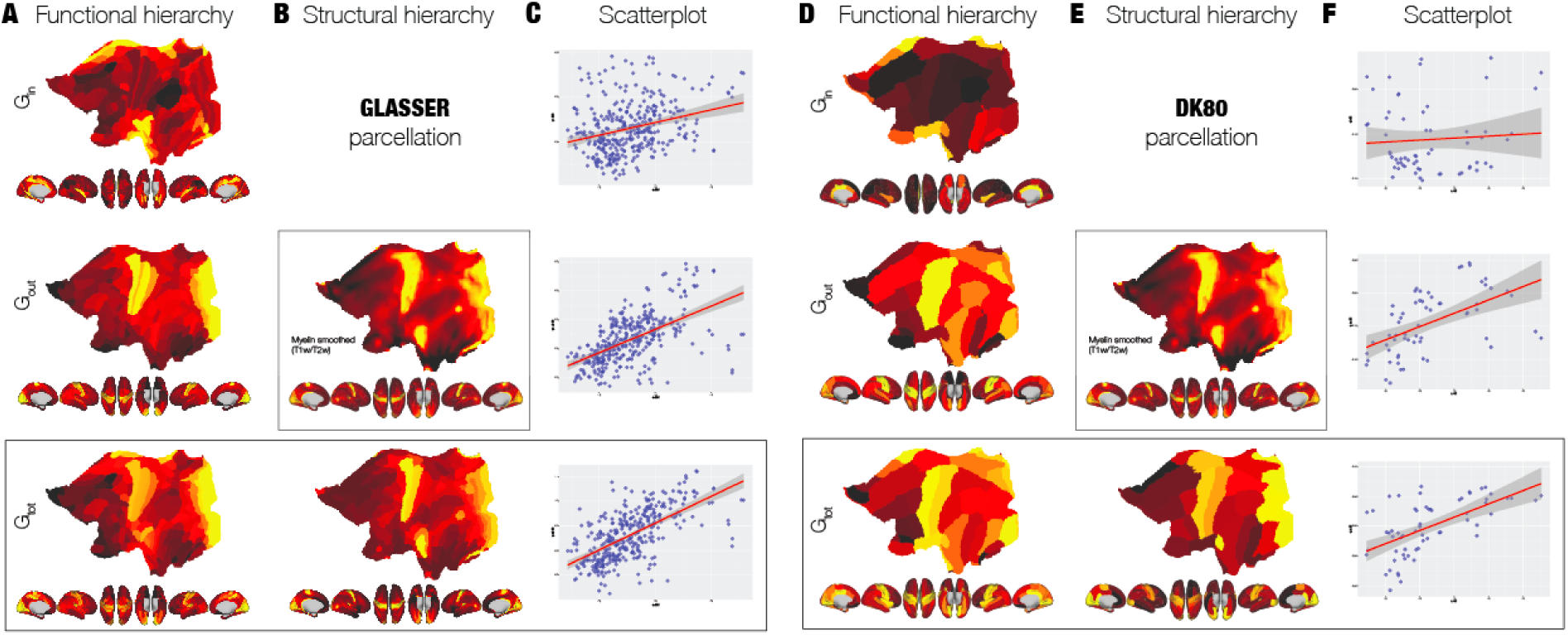
Comparing functional and structural hierarchical organisation. **A)** Functional hierarchy is shown cortical renderings of each of the incoming (G_in_), outgoing (G_out_) and total (G_tot_) directional information flow computed from the NDTE matrix of 1003 HCP participants (see Figure 1 and Methods) in the Glasser parcellation (using renderings of cortical flattening and 3D renderings with midline, right, left, top and bottom views). As can be clearly seen, the outgoing information, G_out_, is highest in sensory areas, while incoming information, G_in_, is highest in higher-order, integrative transmodal areas. **B)** The structural hierarchy is shown for myelination of brain regions (myelin-weighted T1w/T2w). We use the same renderings at the voxel level (top box) and in the Glasser parcellation (bottom). **C)** Scatterplots between the functional hierarchy (G_in_, G_out_ and G_tot_) and structural hierarchy (myelination). The linear correlations are shown by the red line (with standard error in shaded gray) overlaid on the scatterplots. This shows that myelination is remarkably highly correlated with G_tot_, and mainly driven by correlation with the outgoing flow, G_out_,. On the other hand, there is a much lower correlation with the incoming flow, ie integrative measure of G_in_. This means that the static measure of myelination is likely to mostly reflect the driving flow in sensory areas but provides much less information on integrative areas. **D)** Shows the same panel A but for the DK80 parcellation. **E)** Shows the myelination in the DK80 parcellation. **F)** Shows the scatterplots between the functional hierarchy (G_in_, G_out_ and G_tot_) and structural hierarchy (myelination) in the DK80 parcellation. The linear correlations are shown by the red line (with standard error in shaded gray) overlaid on the scatterplots. The results in this coarser scale DK80 parcellation are fully consistent with the finer scale Glasser parcellation.

On the other hand, a popular proxy for anatomical hierarchy is the myelination of brain regions as measured by myelin-weighted T1w/T2w (Glasser and Van Essen, 2011), and so we render this in the Glasser parcellation (**Figures 2B and S1**) and DK80 (**Figure 2E and S1**). Important information about the driving nature of sensory areas (more myelin) and the integrative role of higher-order transmodal areas (less myelin) have been demonstrated from this structural information in recent papers (Burt *et al.*, 2018; Demirtas *et al.*, 2019; Margulies *et al.*, 2016). However, this structural measure does not change with different tasks and is therefore unlikely to capture the functional dynamic changes in hierarchical organisation.

We provide scatterplots between the functional hierarchy (G_in_, G_out_ and G_tot_) and structural hierarchy (myelination) for the cortical regions in the Glasser parcellation (**Figure 2C**) and DK80 (**Figure 2F**). The linear correlations are shown by the red line (with standard error in shaded gray) overlaid on the scatterplots with Glasser values: 0.29 for G_in_, 0.57 for G_out_ and 0.63 for G_tot_ and DK80 values: 0.07 for G_in_, 0.53 for G_out_ and 0.60 for G_tot_. This shows that myelination is remarkably highly correlated with G_tot_, and mainly driven by correlation with the outgoing flow, G_out_. Importantly, as expected, the level of correlation is decreasing between function and structure for the incoming flow (the integrative measure of G_in_).

In order to validate the results from fMRI, we also characterised the functional hierarchical organisation for the corresponding HCP MEG timeseries in the 62 cortical regions of the DK80 parcellation. Attesting to robustness of the results, we found similar correlations between G_in_ and G_out_ from HCP MEG data and myelinisation: 0.04 for G_in_ (14-22Hz, window size 1000ms), 0.48 for G_out_ (22.5-30.5Hz, window size 500ms) (see Supplementary **Figure S2**).

### Functional hierarchy across different tasks and rest

It is clear from these results that the measures of G_in_ and G_out_ are very different and we quantified the changes in the relationship between the functional hierarchy across different tasks and rest. **Figure 3A** provides cortical renderings of all seven tasks and rest of the incoming (G_in_), outgoing (G_out_) and total (G_tot_) directional information flow computed from the NDTE matrix of HCP participants (see **Figure 1** and Methods) in the DK80 parcellation (using 3D views from the side and midline).

**Figure 3.**
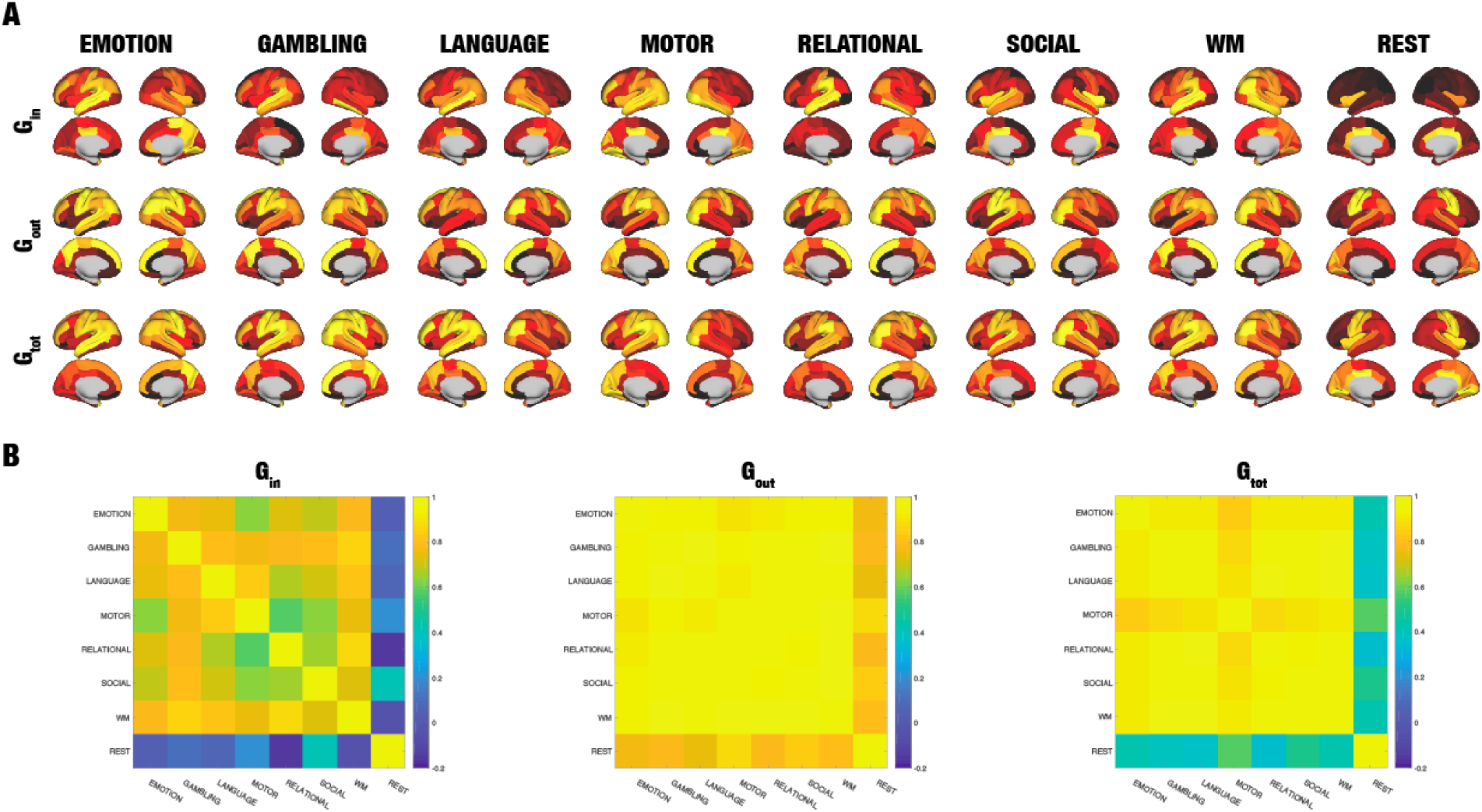
Functional hierarchy across different tasks and rest. **A)** Cortical renderings of all seven tasks and rest of the incoming (G_in_), outgoing (G_out_) and total (G_tot_) directional information flow computed from the NDTE matrix of 1003 HCP participants (see Figure 1 and Methods) in the DK80 parcellation (using 3D views from the side and midline). **B)** Matrices of the comparison of incoming (G_in_), outgoing (G_out_) and total (G_tot_) directional information flow in the seven tasks and rest. As can be clearly seen from G_in_ matrix and the renderings of the incoming flow of information (receivers) are significantly different between tasks and rest. This suggests that different tasks process incoming differently. This is in contrast to the G_out_ matrix and the renderings of the outgoing flow of information (drivers), which are remarkably similar. This shows that sensory areas are consistently driving the information flow. Interestingly, as can be seen from the G_tot_ matrix, the total processing of information flow is more similar within the seven tasks than compared to rest, suggesting the extrinsic, sensory nature of task processing compared to the intrinsic nature of resting state processing.

Quantification of the differences is provided by the correlation matrices between incoming (G_in_), outgoing (G_out_) and total (G_tot_) directional information flow in the seven tasks and rest. The results show that there are significantly different between the hierarchy in tasks and rest for G_in_ (receivers), which reflects that different tasks process incoming information differently. On the other hand, outgoing flow of information (drivers) is remarkably similar across tasks, suggesting that sensory areas are consistently driving the information flow. Importantly, as shown in the G_tot_ matrix, the total processing of information flow is more similar within the seven tasks than compared to rest.

### Quantifying the Functional Rich Club (FRIC) in tasks and resting state

In the structural domain, Zamora-Lopez, Van Heuvel and Sporns proposed the concept of a structural “rich club” (van den Heuvel and Sporns, 2011; Zamora-Lopez *et al.*, 2010), which is characterized by a tendency for high-degree brain regions to be more densely connected among themselves than regions of a lower degree, providing important information on the higher-level topology of the brain network. We extended this concept to the functional domain by defining the concept of a functional rich club (FRIC), which crucially is not static but changes between tasks and rest (see above and Methods).

We computed the NDTE matrices for the DK80 parcellation for all seven tasks (emotion, gambling, language, motor, relational, social, working memory) and resting state for the HCP participants. This allows us to compute the ‘Functional Rich Club’ (FRIC) as the set of regions, separate from the rest of the brain, that define a ‘club’ of functional hubs characterized by a tendency to be more densely functionally connected among themselves than to other brain regions from where they receive integrative information (see Methods). **Figure 4A** shows that the functional rich clubs across tasks and rest contain similar but not identical core regions.

**Figure 4.**
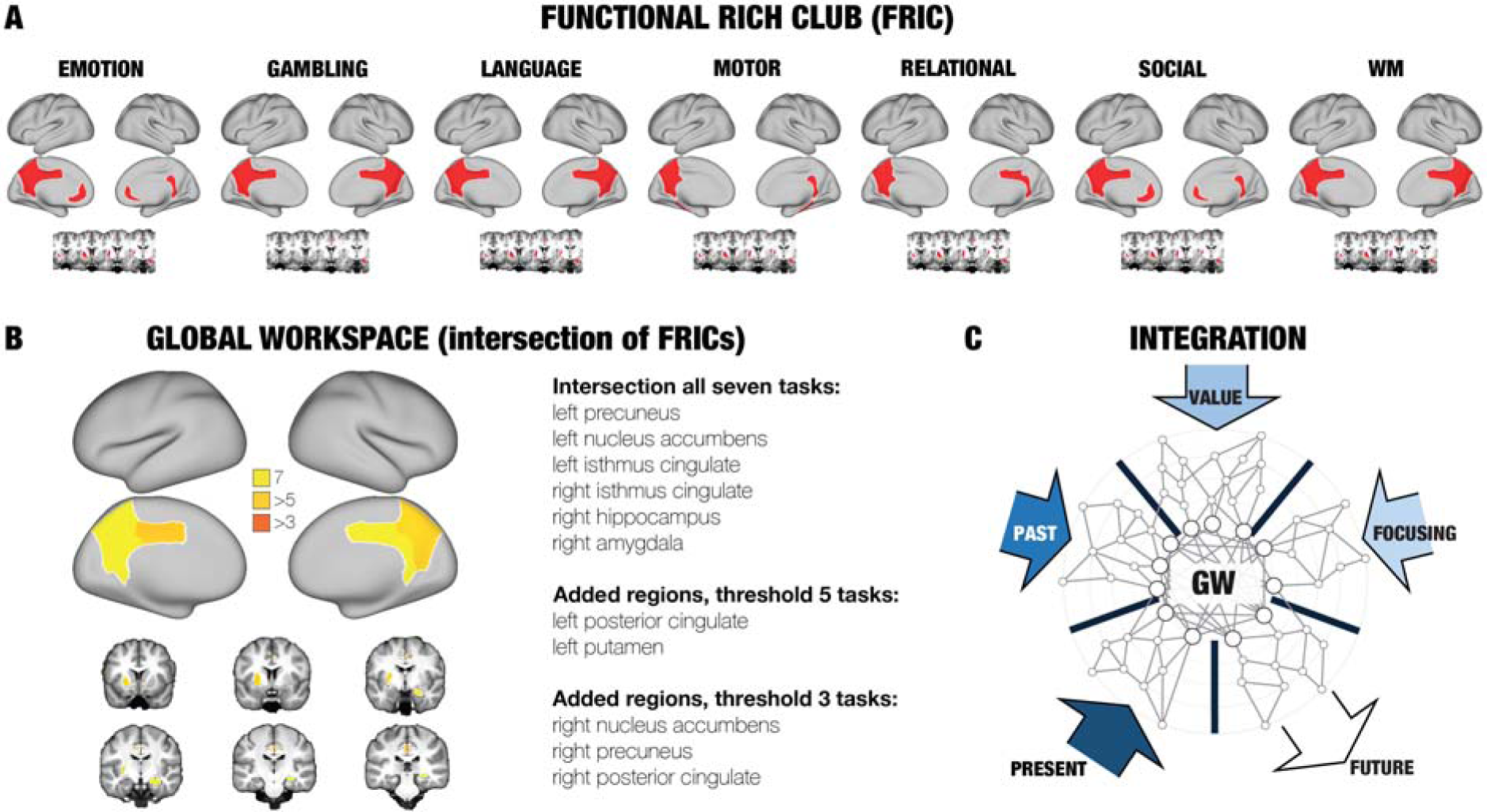
Discovering global workspace as the intersection of functional rich clubs for rest and seven tasks. **A)** We computed the functional hierarchical organisation of all seven tasks (emotion, gambling, language, motor, relational, social, working memory) for the HCP participants. This allows us to compute the ‘Functional Rich Club’ (FRIC) as the set of regions, separate from the rest of the brain, that define a ‘club’ of functional hubs characterized by a tendency to be more densely functionally connected among themselves than to other brain regions from where they receive integrative information (see Methods). As can be seen, these functional rich clubs are similar but not identical across tasks. B) We compute the regions in the Global Workspace (GW) as the intersection of the FRIC members across all possible tasks and resting state. Here, we used the maximal amount of tasks available to provide a reliable estimate of the GW. At the bottom of the figure, we show a rendering of the cortical and subcortical regions in the GNW. As can be seen, the FRIC regions for all seven tasks and rest defining the GNW are the following six brain regions: left precuneus, left nucleus accumbens, right hippocampus, right amygdala and left and right isthmus cingulate. Lowering the threshold of participation to more than five tasks adds two more regions: left posterior cingulate (in five tasks) and left putamen (in six tasks). Further lowering the threshold to three tasks provides another three brain regions: right nucleus accumbens (in three tasks), right precuneus (in three tasks) and right posterior cingulate (in four tasks). C) These regions fit well with the idea suggested by Dehaene and Changeux that the Global Workspace is ideally placed for integrating information from perceptual (PRESENT), long-term memory (PAST), evaluative (VALUE) and attentional (FOCUSING) systems.

### Quantifying the Global Workspace

We quantified the Global Workspace (GW), which is defined by the intersection of FRIC members across all possible tasks and resting state. The HCP data provides resting data as well as seven very different tasks, which we used to provide a reliable estimate of the GNW. **Figure 4B** plots a rendering of the cortical and subcortical regions in the GNW.

We found that the intersecting FRIC members for all seven tasks and rest are the following six brain regions: left precuneus, left nucleus accumbens, right hippocampus, right amygdala, left and right isthmus cingulate. Searching for a less restrictive definition, we further lowered the threshold to include areas only common to six tasks and rest, adding the left putamen and further lowering to five tasks, which added the left posterior cingulate. Even more we found that lowering the threshold to four tasks added the right posterior cingulate, while common to three tasks added right nucleus accumbens and right precuneus. The results point to a remarkable stable, non-lateralised core of brain regions necessary in the global workspace.

### Establishing causal significance of FRIC by lesioning whole-brain model

Still, this quantification is not causal and we wanted to establish the causal mechanistic functional relevance of the FRIC brain regions orchestrating functional hierarchical organisation of the resting state. So we created a whole-brain model that can generate the effective connectivity underlying the functional hierarchy and subsequently be manipulated (see Methods). **Figure 5A** shows the general framework of how this model uses optimised anatomically-constrained parameters (generative anatomically-constrained bidirectional connectivity, GABIC) to describe the effective strength of the synaptic coupling to fit the NDTE matrix by maximising the empirical and simulated NDTE matrices. The NDTE matrix is a strong measure of the effective connectivity in the brain and consequently GABIC is a matrix *generating* this effective connectivity rather than the effective connectivity *per se*. Optimising the model using a particle swarm optimizer (see Methods) was computationally very demanding but we managed to find a very good fit with a correlation between empirical and model-generated NDTE matrices of 0.6. Please note that given that NDTE matrix is bidirectional and the GABIC matrix is therefore also asymmetric. Importantly, the GABIC matrix does not correlate with the NDTE matrix (correlation of only 0.01), demonstrating the complex non-linear relationship between the generative parameters and the effective connectivity as measured by NDTE. NDTE correlated 0.49 with the static functional connectivity, demonstrating that there is complementary information in the causal information flow measure of NDTE thus further constraining the whole-brain modelling.

**Figure 5.**
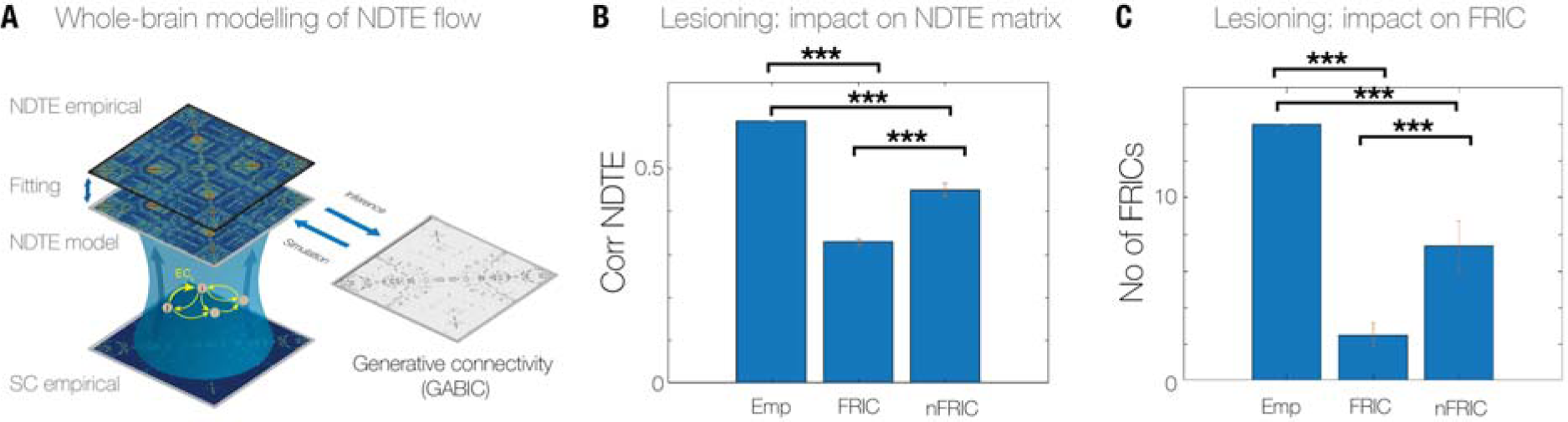
Establishing causal significance by lesioning whole-brain model. **A)** In order to establish the causal mechanistic functional relevance of the FRIC brain regions orchestrating functional hierarchical organisation of the resting state, we created a whole-brain model to fit the empirical NDTE flow matrix. **B)** Lesioning all FRIC compared to non-FRIC members in this whole-brain model causes significant breakdown in information flow. The correlation between empirical and simulated NDTE matrices is maximally affected when FRIC members are lesioned (p<10^-30^) compared to when non-FRIC members are lesioned. **C)** Similarly, we found significant differences in the number of FRIC members detected when lesioning the original empirical FRIC members compared to empirical non-FRIC members (p<10^-30^). This is causal evidence for the major significance of FRIC in orchestrating functional organisation.

Here, we used the whole-brain model to ascertain the causal significance of the FRIC regions in resting state. **Figure 5B** shows the result of lesioning all FRIC compared to non-FRIC members in the whole-brain model, which causes very significant breakdown in information flow. When measuring this in terms of correlation between empirical and simulated NDTE matrices we found that this impacts highly significantly the fitting when FRIC members are lesioned (p<10^-30^) compared to the lesioning of non-FRIC members. Furthermore, in **Figure 5C**, we show the significant consequences of detecting significant differences in the number of FRIC members detected when lesioning the original empirical FRIC members compared to empirical non-FRIC members (p<10^-30^). This is strong causal evidence for the major significance of FRIC in orchestrating functional organisation.

### Alternative measure of functional hierarchy

Finally, we were inspired by the careful neuroanatomical research by Markov and colleagues to define a similar measure of feedforward and feedback organisation but here for the functional domain (Felleman and Van Essen, 1991; Markov *et al.*, 2013; Markov *et al.*, 2014) (see Methods). However, as can be seen in **Figure S3**, this FF Hierarchy measure is only weakly discriminatory between the seven tasks – and only weakly correlated with G_in_, the integrative measure of incoming information flow.

## Discussion

The major advance over the last couple of years in acquiring large multimodal neuroimaging datasets (‘big data’) is finally making it possible to address large, challenging problems in systems neuroscience (Alfaro-Almagro *et al.*, 2018; Glasser *et al.*, 2016b; Poldrack and Gorgolewski, 2014). A key open problem is how best to characterise the functional hierarchical organisation of whole-brain dynamics to understand the orchestration of brain processing. Here, we applied our novel *Normalised Directed Transfer Entropy* (NDTE) framework to over 1000 participants from the Human Connectome Project (HCP) (Van Essen *et al.*, 2013). This provided the precise characterisation of the causal flow of information between pairs of brain regions and consequently the estimation of the overall hierarchical organisation of the human brain in resting state and across seven tasks. This revealed the functional rich club (FRIC) organisation of a core group of brain regions changing with task and rest. Crucially, this advance allowed us to discover the core regions in the ‘global workspace’ (Baars, 1989; Dehaene *et al.*, 1998) as the common FRIC regions invariant across task and rest.

### Discovering the Global Workspace

The global workspace was found to consist of a core subset of brain regions including the precuneus, posterior and isthmus cingulate, nucleus accumbens, putamen, hippocampus and amygdala. This core functional ‘club’ of integrative brain regions is remarkably consistent with the original proposal by Dehaene and Changeux (Dehaene *et al.*, 1998), which suggests that the global neuronal workspace must integrate past and present through focusing and evaluation. Indeed, the authors propose that associative perceptual, motor, attention, memory, and value areas interconnect to form a higher-level unified space. For the integration of the past, the hippocampus has been shown to play a key role in many aspects of memory (see for example (Eichenbaum *et al.*, 2007; Scoville and Milner, 1957; Squire *et al.*, 2004)). Similarly, the evaluation of value has been shown to involve the nucleus accumbens (e.g. (Berridge and Kringelbach, 2008; Berridge and Robinson, 2003; Haber and Knutson, 2010)), putamen (e.g. (Di Martino *et al.*, 2008; Haber and Knutson, 2010)) and the amygdala (e.g. (Haber and Knutson, 2010; LeDoux, 2003; Schoenbaum *et al.*, 2003; Swanson and Petrovich, 1998; Zald and Pardo, 1997)). The integration of the past, present and future by processing and attending perceptual information has been strongly associated with the precuneus (e.g. (Cavanna and Trimble, 2006; Margulies *et al.*, 2009; Northoff and Bermpohl, 2004)) and the posterior and isthmus cingulate cortices (e.g. (Haber and Knutson, 2010; Leech and Sharp, 2014; Mesulam, 1999; Northoff and Bermpohl, 2004; Paus, 2001)). Interestingly, the functions of the precuneus have also been shown to be compromised in coma and vegetative state (Laureys *et al.*, 2004).

The definition of the global workspace proposed here follows the original ideas of Baars’ cognitive theory of consciousness, which distinguishes a vast array of unconscious specialised processors running in parallel, and a single limited-capacity serial “global workspace” that allows them to exchange information (Baars, 1989). The subsequent development by Dehaene and Changeux (Dehaene and Changeux, 2005) of this theory into the *global neuronal workspace* includes a further “ignition” component capturing the strong temporary increase in synchronized firing leading to a coherent state of activity. The transition to this state of high correlated activity is very fast and leads to amplification of local neural activation and the subsequent *ignition* of multiple distant areas. We have not studied ignition here since the fast timescale (typically about 100-200 ms) is difficult to capture with fMRI – although we have recently shown that such fast timescales can be reconstructed using appropriate whole-brain modelling (Deco *et al.*, 2019b). Yet, the potential role of ignition in initiating and sustaining FRIC and the global workspace should clearly be further investigated in future studies using for example MEG.

### Causal evidence for importance of FRIC regions

Importantly, as a further proof of causal significance of the core FRIC regions, we built and lesioned a whole-brain model that can generate effective connectivity obtained using our NDTE framework (see **Figure 5** and Methods). We found that lesioning ten FRIC regions for the resting state very significantly impaired the flow of information and the ability to form new FRIC among the remaining brain regions. This causally establishes the full FRIC network as having a key integrative role in orchestrating functional hierarchical organisation. Furthermore, we found that this FRIC network is fairly fault tolerant in that lesioning four FRIC members were not significantly different from lesioning four non-FRIC members. However, lesioning five FRIC compared to five non-FRIC members did lead to significantly different information flow. This suggests that there is a tipping point where the breakdown of information flow in the FRIC leads to significant problems. As such, the partial breakdown of FRIC members could be a significant factor in the transitioning to neuropsychiatric disorders (Deco and Kringelbach, 2014). In particular the specific breakdown of the integration of the evaluative system related to self-processing (precuneus, nucleus accumbens, putamen and cingulate cortices) and could lead to the anhedonia, the lack of pleasure, which is a major symptom of neuropsychiatric disease (Berridge and Kringelbach, 2015; Kringelbach, 2005).

### Implications for elucidating functional brain hierarchy

A long history of neuroanatomical discoveries has demonstrated that the brain is clearly hierarchical in its structure from single units to the larger circuits (Bullmore and Sporns, 2012; Felleman and Van Essen, 1991; Hagmann *et al.*, 2008; Markov *et al.*, 2014; Mesulam, 1998; van den Heuvel and Sporns, 2011; Zamora-Lopez *et al.*, 2010). Research by Margulies and colleagues (Margulies *et al.*, 2016) have used neuroimaging to extend Mesulam’s seminal proposal that brain processing is shaped by a hierarchy of distinct unimodal areas to integrative transmodal areas (Mesulam, 1998). Recently, this idea has been further extended by applying the principle of harmonic modes to functional connectivity HCP data (Glomb *et al.*, 2019). This revealed hitherto unknown principle unifying the gradiental and modular aspects and revealing the multi-dimensional hierarchical nature of brain organisation.

Here, we have developed the novel NDTE framework for discovering the functional hierarchical organisation of the human brain. We have demonstrated that this can be used to characterise the FRIC corresponding to the core integrative transmodal brain regions allowing for the necessary whole-brain cohesion (as shown by the cartoon in a simplified 2D representation in **Figure 1C**). As shown in **Figure 2**, the outgoing information flow (drivers), *G*_*out*_, reflects mostly the unimodal sensory regions, where the incoming information flow (integrative receivers), *G*_*in*_, reflects the higher-order transmodal regions.

The results using the NDTE framework confirm and extend previous neuroanatomical findings of the static anatomical hierarchy measuring through cortical myelination and in particular the neuroimaging measure of weighted T1w/T2w maps (Glasser and Van Essen, 2011) as a useful proxy for hierarchy (Burt *et al.*, 2018; Demirtas *et al.*, 2019; Margulies *et al.*, 2016). The results show that the NDTE framework can capture this static myelination hierarchy (**Figure 2 and S1**). Specifically, we found that the sum of incoming and outgoing flow of information, *G*_*tot*_, correlates very highly with the T1w/T2w maps in both the Glasser and DK80 parcellations. Using causal measures of timeseries have had a long history and there have been considerable arguments in the literature as to the appropriateness of using these on BOLD timeseries due to the potential impact of the variability of the haemodynamic response across brain regions (David *et al.*, 2008; Friston, 2009; Smith *et al.*, 2011). Nevertheless, it has also been demonstrated that causal timeseries methods perform much better with sufficiently fast sampling and low measurement noise (Seth *et al.*, 2013). This is the case with the state-of-the-art HCP data with a fast TR of 0.78 seconds, providing excellent subsampling of the haemodynamic response function, and thus mitigating the potential problems with poor subsampling with long TR that can lead to problems with spurious and undetectable causality, as well as distortion of relative strengths.

Still, we also confirmed the results by applying the NDTE framework to MEG timeseries from the 62 cortical regions of the DK80 parcellation, and demonstrating similarly high correlations with myelination (**Figure S2**). This clearly demonstrates the robustness of the NDTE framework for both haemodynamic and direct electromagnetic measures of brain activity.

The NDTE framework includes suitable normalisation, surrogates and p-value aggregation across large number of participants (see Methods), which contribute to robustness of the method. Indeed, the high correlation between myelination and the total incoming and outgoing information flow, *G*_*tot*_, obtained for both BOLD and MEG, provides strong empirical evidence and confidence in the highly meaningful results.

Complementary to the measures of causal connection strength, we also designed another hierarchical measurement inspired by the seminal GLM method of computing the fraction of feedforward and feedback organisation originally developed in neuroanatomy (Felleman and Van Essen, 1991; Markov *et al.*, 2013; Markov *et al.*, 2014). **Figure S3** shows that this FF Hierarchy measure is highly correlated with itself across the seven tasks and rests. In contrast this measure correlates only very weakly with *G*_*in*_, the integrative measure of incoming information flow. This is expected given that FF hierarchy is an ordered measure of the fraction of feedforward and feedback connections. As such this is not as useful for establishing the full hierarchical organisation as the other NDTE flow measures.

### Causal confirmation using novel generative whole-brain model

Traditionally, whole-brain models have been relatively successful in linking structural connectivity with functional dynamics (Breakspear, 2017; Deco and Kringelbach, 2014). This has revealed important new mechanistic principles of brain function (Deco *et al.*, 2018; Deco *et al.*, 2019a; Deco *et al.*, 2019b; Deco *et al.*, 2017e; Honey *et al.*, 2007). Nevertheless, the present causal characterisation of whole-brain information flow offers a new avenue for generating even more useful models. We have created a generative whole-brain model that can recreate the causal information flow in terms of the NDTE matrix. In other words, our second order model using GABIC to describe the generators accounting for the causal influence of one neural system over another.

Crucially, this whole-brain model was used to systematically test the robustness and fault tolerance of the empirically extracted FRIC members. Lesioning all FRIC compared to non-FRIC members causes very significant breakdown in information flow (p<10^-30^). This provides causal mechanistic evidence for the importance of the FRIC members in orchestrating the complex choreography of information flow within the functional hierarchical organisation of the human brain.

### Conclusion

The findings presented here help shed new light on a major unsolved problem in neuroscience which is to characterise the functional hierarchical organisation of whole-brain dynamics underlying the orchestration of information across the whole-brain. Crucially, this enabled us to discover the core regions of the global workspace across task and rest. The brain regions identified corresponds remarkably well to the predictions made by Changeux and Dehaene of higher-level unified space which is well suited for integrating past and present through focusing and evaluation. While the results presented here pertain only to the global workspace of conscious processing, future work could use the NDTE framework to investigate other states such as sleep and anaesthesia. Such further investigations could potentially allow for a direct comparison between the global workspace theory with other theories of consciousness such as the Integrated Information Theory (Tononi *et al.*, 1994) and the Temporo-spatial Theory of Consciousness (Northoff, 2013). Equally, the NDTE framework could be used to investigate unbalanced brain states in neuropsychiatric disorders. Importantly, given that such investigations would be using a generative whole-brain model, this could subsequently be used to perturb and rebalance the model to discover novel optimal, causal paths to health (Deco *et al.*, 2017a; Deco *et al.*, 2019a; Deco and Kringelbach, 2014).

## Methods

### Neuroimaging data acquisition, preprocessing and timeseries extraction

#### Ethics

The Washington University–University of Minnesota (WU-Minn HCP) Consortium obtained full informed consent from all participants, and research procedures and ethical guidelines were followed in accordance with Washington University institutional review board approval.

#### Participants

The data set used for this investigation was selected from the March 2017 public data release from the Human Connectome Project (HCP) where we chose a sample of 1003 participants. From this large sample we further chose to replicate in the smaller subsample of 100 unrelated participants (54 females, 46 males, mean age □=□29.1□+/-□3.7 years). This subset of participants provided by HCP ensures that they are not family relatives, and this criterion was important to exclude possible identifiability confounds and the need for family-structure co-variables in the analyses.

#### Neuroimaging acquisition for fMRI HCP

The 1003 HCP participants were scanned on a 3-T connectome-Skyra scanner (Siemens). We used one resting state fMRI acquisition of approximately 15 minutes acquired on the same day, with eyes open with relaxed fixation on a projected bright cross-hair on a dark background as well as data from the seven tasks. The HCP website (http://www.humanconnectome.org/) provides the full details of participants, the acquisition protocol and preprocessing of the data for both resting state and the seven tasks. Below we have briefly summarised these.

#### Neuroimaging acquisition for dMRI HCP

Diffusion spectrum and T2-weighted imaging data from 32 participants in the Human Connectome Project (HCP) were obtained from the HCP database. The acquisition parameters are described in details on the HCP website (Setsompop *et al.*, 2013).

#### Neuroimaging acquisition for MEG HCP

We used the human non-invasive resting state magnetoencephalography (MEG) data publicly available from the Human Connectome Project (HCP) consortium, acquired on a Magnes 3600 MEG (4D NeuroImaging, San Diego, USA) with 248 magnetometers. The resting state data consist of 89 subjects (mean 28.7 years, range 22-35, 41 f / 48 m, acquired in 3 subsequent sessions, lasting 6 minutes each).

#### The HCP task battery of seven tasks

The HCP task battery consists of seven tasks: working memory, motor, gambling, language, social, emotional, relational, which are described in details on the HCP website (Barch *et al.*, 2013). HCP participants performed all tasks in two separate sessions (first session: working memory, gambling and motor; second session: language, social cognition, relational processing and emotion processing).

### Neuroimaging structural connectivity and extraction of functional timeseries

#### Parcellations

All neuroimaging data was processed using two standard cortical parcellations with added subcortical regions. For a fine-scale parcellation, we used the Glasser parcellation with 360 cortical regions (180 regions in each hemisphere) (Glasser *et al.*, 2016a). We added the 18 subcortical regions, ie nine regions per hemisphere: hippocampus, amygdala, subthalamic nucleus (STN), globus pallidus internal segment (GPi), globus pallidus external segment (GPe), putamen, caudate, nucleus accumbens and thalamus. This created a parcellation with 378 regions: Glasser378 parcellation, which is defined in the common HCP CIFTI grayordinates standard space with a total of 91,282 grayordinates (sampled at 2 mm^3^).

For a coarser-scale parcellation, we used the Mindboggle-modified Desikan-Killiany parcellation (Desikan *et al.*, 2006) with a total of 62 cortical regions (31 regions per hemisphere) (Klein and Tourville, 2012). We added the same 18 subcortical regions mentioned above (9 regions per hemisphere) and ended up with 80 regions in the DK80 parcellation; also precisely defined in the common HCP CIFTI grayordinates standard space.

#### Generating structural connectivity matrices from dMRI

We used the state-of-the-art preprocessed dMRI data from 32 HCP participants available as part of the freely available Lead-DBS software package (http://www.lead-dbs.org/). The precise preprocessing is described in details in Horn and colleagues (Horn *et al.*, 2017) but briefly, the data was processed using a generalized q-sampling imaging algorithm implemented in DSI studio (http://dsi-studio.labsolver.org). Segmentation of the T2-weighted anatomical images produced a white-matter mask and co-registering the images to the b0 image of the diffusion data using SPM12. In each HCP participant, 200,000 fibres were sampled within the white-matter mask. Fibres were transformed into MNI space using Lead-DBS (Horn and Blankenburg, 2016). We subsequently used the standardized methods in Lead-DBS to produce the structural connectomes for both the Glasser378 and DK80 parcellations

#### Preprocessing and extraction of functional timeseries in fMRI resting state and task data

The preprocessing of the HCP resting state and task datasets is described in details on the HCP website. Briefly, the data is preprocessed using the HCP pipeline which is using standardized methods using FSL (FMRIB Software Library), FreeSurfer, and the Connectome Workbench software (Glasser *et al.*, 2013; Smith *et al.*, 2013). This preprocessing included correction for spatial and gradient distortions and head motion, intensity normalization and bias field removal, registration to the T1 weighted structural image, transformation to the 2mm Montreal Neurological Institute (MNI) space, and using the FIX artefact removal procedure (Navarro Schroder *et al.*, 2015; Smith *et al.*, 2013). The head motion parameters were regressed out and structured artefacts were removed by ICA+FIX processing (Independent Component Analysis followed by FMRIB’s ICA-based X-noiseifier (Griffanti *et al.*, 2014; Salimi-Khorshidi *et al.*, 2014)). Preprocessed timeseries of all grayordinates are in HCP CIFTI grayordinates standard space and available in the surface-based CIFTI file for each participants for resting state and each of the seven tasks.

We used a custom-made Matlab script using the ft_read_cifti function (Fieldtrip toolbox (Oostenveld *et al.*, 2011)) to extract the average timeseries of all the grayordinates in each region of the Glasser and DK80 parcellations, which are defined in the HCP CIFTI grayordinates standard space.

#### Preprocessing and extraction of MEG data timeseries

For each participant, the MEG data were acquired in a single continuous run comprising resting state. As a starting point we used the ‘preprocessed’ MEG data from the HCP database. At this level of preprocessing, removal of artefactual independent components, bad samples and channels have already been performed (Larson-Prior *et al.*, 2013). We then subjected the preprocessed data to bandpass filtering (1-48Hz, Butterworth) and LCMV beamforming (using beamforming routines from the Matlab based Fieldtrip toolbox (Oostenveld *et al.*, 2011)), which is downsampled to 200 Hz and resulted in 5798 virtual source voxels (with 8mm grid resolution). We extracted timecourses from the 62 cortical regions of the DK80 parcellation. All resting state runs for a participant were acquired in a single session and we concatenated the resting state runs for each participant, and applied a single beamformer, parcel time-course extraction and spatial leakage reduction.

### Neuroimaging analysis tools and methods

#### Normalized Directed Transfer Entropy (NDTE)

In order to establish and to investigate the *functional* hierarchical organisation of whole-brain activity we need first to characterize how different brain regions communicate between each other, i.e. compute the directed flow between regions. We characterize the functional interaction between two brain regions, in a given parcellation, by an information theoretical statistical criterion that allows us to infer the underlying bi-directional causal reciprocal communication. Let us assume that we want to describe the statistical causal interaction exerted from a source brain area *X* to another target brain area *Y*. We aim to measure the extra knowledge that the dynamical functional activity of the past of *X* contribute to the prediction of the future of *Y*, by the following mutual information:

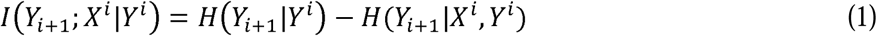

where *Y*_*i+1*_ is the activity level of brain area *Y* at the time point *i+1*, and *X*^*i*^ indicates the whole activity level of the past of *X* in a time window of length *T* up to and including the time point *i* (i.e. *X*^*i*^*=[X*_*i*_ *X*_*i-1*_ … *X*_*i-(T-1)*_]). Note that this causality measure is not symmetric, i.e. allows bidirectional analysis. The conditional entropies are defined as follows:

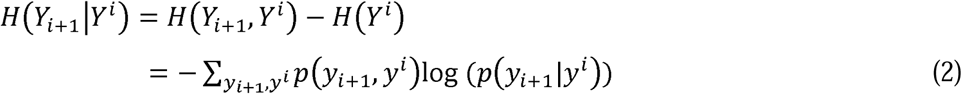

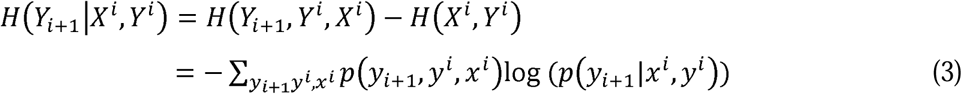

The mutual information *I(Y*_*i+1*_; *X*^*i*^*|Y*^*i*^) expresses the degree of statistical dependency between the past of *X* and the future of *Y*. In other words, if that mutual information is equal to zero, then the probability *p*(*Y*_*i+1*_; *X*^*i*^*|Y*^*i*^) *= p*(*Y*_*i+1*_*|Y*^*i*^)· *p*(*Y*_*i+1*_*|Y*^*i*^) and thus we can say that there is no causal interaction from *X* to *Y*.

Consequently *I(Y*_*i+1*_; *X*^*i*^*|Y*^*i*^) expresses a strong form of Granger causality (Granger, 1980), by comparing the uncertainty in *Y*_*i+1*_ when using knowledge of only its own past *Y*^*i*^ or the past of both brain regions, i.e. *X*^*i*^, *Y*^*i*^. This information-theoretical concept of causality was introduced in neuroscience by Schreiber (2000) and is usually called *Transfer Entropy* (Brovelli *et al.*, 2015; Chicharro and Ledberg, 2012; Vicente *et al.*, 2011; Wibral *et al.*, 2014). In order to facilitate computation, Brovelli et al. (2015) proposed a weaker form of causality allowing calculation of the involved entropies by just considering a Gaussian approximation, i.e by considering only second order statistics. Indeed, under this approximation, the entropies can be computed as follows:

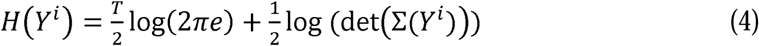

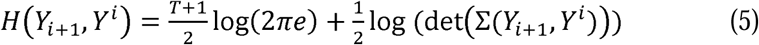

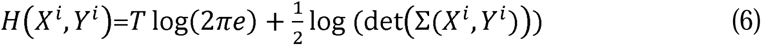

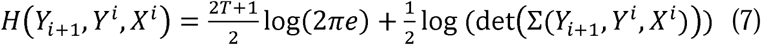

In other words, causality is based only on the corresponding covariance matrices.

In order to be able to sum and compare the directed mutual information flow between different pairs of brain regions, this has to be appropriately normalised. In fact, if the mutual information directed flow is correctly normalized then the different values could be combined for example to know the total directed flow exerted by the whole brain on a single region or vice versa, the directed flow exerted by a single brain region on the whole brain. More specifically, we define this information theoretical measure as normalised directed transfer entropy (NDTE) flow:

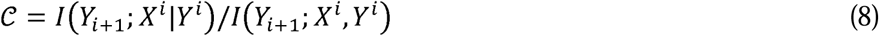

where *I(Y*_*i+1*_; *X*^*i*^*|Y*^*i*^) is the mutual information that the past of both signal together, *X*^*i*,^,*Y*^*i*^, has about the future of the target brain region *Y*_*i+1*_. Given that,

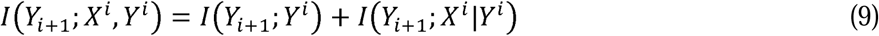

this normalisation compares the original mutual information directed flow, i.e. the predictability of *Y*_*i+1*_ by the past of *X*^*i*^*|Y*^*i*^ with the internal predictability of *Y*_*i+1*_, i.e. *I(Y*_*i+*1_; *Y*^*i*^).

Furthermore, in order to perform a solid and robust statistical significance analysis of the NDTE flow, 𝒞, we use the surrogate framework, inspired by the work of Theiler and colleagues (Theiler *et al.*, 1992). This traditional surrogate methodology uses a phase randomisation of the Fourier transform of the original data in order to preserve the linear correlations. Nevertheless, as discussed and analysed rigorously in Diks and Fang (Diks and Fang, 2017); these methods are not suitable for detecting significance when using entropy measures, as discussed extensively (Faes *et al.*, 2008; Hinich *et al.*, 2005). In view of these problems, Quiroga and colleagues proposed the circular time shifted surrogates method, which is a robust method using surrogates that can be used for causality measurements (Quiroga *et al.*, 2002). Hence, we use this methodology for analysing the p-values of the hypothesis testing, aiming to detect significant values in 𝒞 for each pair for each single participant. For each statistical test (i.e. each pair of regions and each subject), we generate 100 independent circular time-shifted surrogates by separately resampling both the driving signal *X* and the target response signal *Y*. Specifically, two independent random integers *c* and *d* are randomly generated within the interval [0.05n 0.95n] (where *n* is the number of time point in the time series signal). Then the circular time-shift is performed by moving the first *c* values of *X*=[*X*_1_, …, *X*_*n*_] to the end of the time series which creates the surrogate sample *X*=[*X*_*c* + 1_, …, *X*_*n*_, *X*_1_, …, *X*_*c*_], and similarly for *Y* the first *d* values of *Y*=[*Y*_1_, …, *Y*_*n*_]are moved to the end of the time series to create the surrogate sample *Y*=[*Y*_*d* + 1_, …, *Y*_*n*_, *Y*_1_, …, *Y*_*d*_].

This type of surrogates does not assume Gaussianity and preserves the whole statistical structure of the original time series. We use a nonparametric kernel distribution representation of the probability density function of the surrogate values of 𝒞, and compare the fraction of area of that distribution above the value of the NDTE flow of the original data, 𝒞_*original*_, to the total area, and compute the corresponding *p*-value.

After computing the individual *p*-values for each brain region pairs and each single participant, we aggregate the *p*-values for each single pair of brain areas across the whole group of participants. The combination of different *p*-values across subjects is a classical problem in statistics that was originally addressed by Ronald Fisher in what nowadays is known as Fisher’s method (Fisher, 1925). Here, we used a more sensitive methodology, namely the Stouffer’s method (Stouffer *et al.*, 1949) which sums the inverse normal transformed *p*-values. Indeed, the Stouffer’s statistics is given by

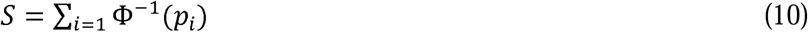

where Φ is the standard normal cumulative distribution function, and *p*_i_ the *p*-values of each participant *i* (computed for a given pair). Under the null hypothesis, the Stouffer’s statistics is normal distributed *N(0,m)*, being *m* the total number of participants. After the aggregation of the pairs of *p*-values across participants, we correct for multiple comparisons by using the traditional False Discover Rate method of Benjamini and Hochberg (Benjamini and Hochberg, 1995).

The result of the significance test across participants determines a binary matrix *T* (with dimension number of brain regions in a given parcellation) that indicates with ones or zeros if the corresponding pair is significant or not (rows indicates target regions and columns driving regions). In fact, we can define now with this matrix *T*, the *broadness* of incoming or outgoing information for each brain area *i* by *B*_*in*_(*i*) = ∑_*j*_ *T*_*ij*_, and *B*_*out*_(*i*) = ∑_*j*_ *T*_*ji*_, respectively. The broadness of the incoming or outgoing information is the number of areas that drives or that are influenced significantly across participants by a single specific brain region.

#### Functional Hierarchical Organisation

Using the above methods describing NDTE flow, we can establish and study the functional organisation of data where the different levels of directed flow to and from a given brain region *I* are given *G*_*in*_(*i*), *G*_*out*_(*i*), and *G*_*tot*_(*i*).

Our analysis of functional relevance and hierarchy is based on the resulting averaged NDTE flow, 𝒞_*All*_, across participants. We define for each brain region *i* the incoming level of directed flow, i.e. the degree of being a receiver, by 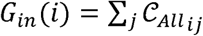. Similarly, for each brain region *i* the outgoing level of NDTE flow, i.e. the degree of being a driving region, by 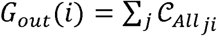. The total level of functional interaction for each brain region *i* is thus given by *G*_*out*_(*i*) = *G*_*in*_(*i*) + *G*_*out*_(*i*).

#### Functional Rich Club (FRIC)

We define the functional rich club (FRIC) in a matrix 𝒞_*All*_ with *N* regions, by running a simple algorithm that searches for the largest subset of regions *k* = [*i*_1_,..,*i*_*l*_], where *G*_*FRIC*_ (*k*) is significantly larger than all other sets with the same number of regions:

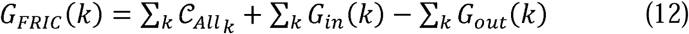

Using a more detailed notation, the equation can be further expressed as

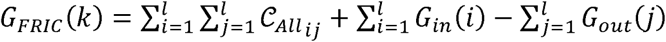

where *l* is the number of regions that defines the subset of regions *k* = [*i*_1_,..,*i*_*l*_].

The first and third term can be transformed to

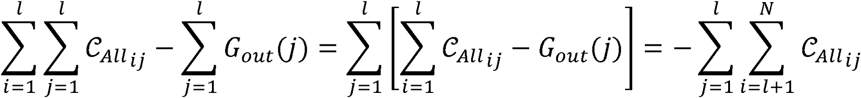

where *N* is the total number of regions.

Similarly, the second term, i.e. the sum of *G*_*in*_, can be expressed as

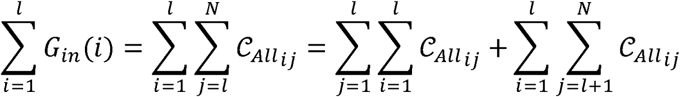

Thus, taken together, Equation 12 can also be written as

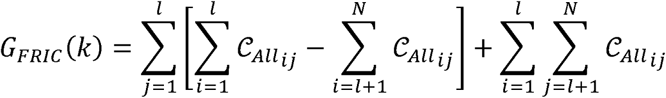

Where the first term describes the difference between the information flow of each member of the subset to the other members in the subset compared with the information flow *outside* the subset. The second term describes the information flow that each member of the subset receives from outside.

As can be appreciated from the combinatorics, for a matrix 𝒞_*All*_ with more than just a few regions, it is computationally very demanding to exhaustively compute the optimal solution. However, it is also clear from the definition that the FRIC for a given set of regions are likely to be found from the regions with highest level of incoming directed flow (*G*_*in*_). So we created the following algorithm: Sort the regions according to *G*_*in*_ and then iteratively computed *G*_*FRIC*_ for progressively more *l* regions. Statistical significance was computed by replacing a random region of the *k* regions with any of the remaining regions using a Monte Carlo framework.

#### Defining hierarchy using GLM

In the literature, hierarchy has traditionally been proposed using neuroanatomy and specifically the anatomical hierarchical ordering based on the different structure of feed-forward and feedback connections across the brain (Barone *et al.*, 2000; Felleman and Van Essen, 1991). Based on this, Markov and colleagues defined a framework for assigning hierarchical values to each region in such a way that the difference of the hierarchical values in two brain regions predicts the fraction of feed-forward connections coupling those two brain regions (Markov *et al.*, 2013; Markov *et al.*, 2014). In fact, they proposed to use the Supragranular Layer Neurons (SLN) index defined as the fraction of projections originated in the supragranular layer of the source area to the target area divided by the total number of projections between the SLN of projections. This idea was based on the observations of Felleman and Van Essen (1991) and Barone et al. (2000) that in the visual system, feed-forward projections directed from early visual areas to higher-order areas tend to originate in the supragranular layers of the cortex and terminate in layer 4, whereas, projections from higher-order areas to early visual areas originate in the infragranular layers and terminate outside of layer 4.

Our framework of computing the NDTE flow (see above) allows us to extend these seminal ideas to the functional level. Instead of using the anatomically based SLN index, we can use the fraction of functional feed-forward causal connections with respect to the total number of connections, feed-forward and feedback, between two brain regions. Consequently, we assign a functional hierarchical value *H* to each brain region such that the difference of the corresponding values between two brain regions predicts the functional fraction of direct connections from one source region *i* to another target region *j*. We use a generalized linear model to establish this prediction, as follows:

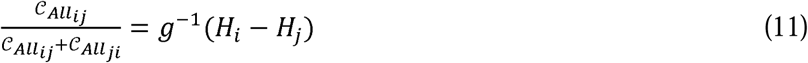

where the left hand is the fraction of feedforward connections respect to the total, g^-1^ is a logistic regression function (which correspond to fitting a Generalized Linear Model (GLM) with a binomial family (McCullagh and Nelder, 1989), and *H*_*i*_ and *H*_*j*_ are the functional hierarchical values inferred for brain regions *i* and *j*. In order to establish a reference point, we assigned a hierarchy value of zero to the last parcel.

#### Whole-brain model of NDTE flow

The main aim of whole-brain modelling is to infer the causal dynamical mechanisms generating the observed empirical spatiotemporal dynamics. Here, we would like to estimate the generators underlying the empirical spatiotemporal dynamics in terms of the causal relationships between the different brain regions, i.e. the NDTE flow. Specifically, the whole-brain model will link the structural anatomy (given by the dMRI data) with the functional dynamics (given by the fMRI data) by adapting the free parameters, i.e. the generators to provide the optimal fit between the simulated and empirical NDTE flow. The generators are internal parameters describing the local dynamics of a brain region, such as noise and latency, as well as the strength of the synaptic conductivity of connections between different brain areas linked by the anatomical fibres. We call the matrix of the generative conductivities of the existing anatomical fibres for “Generative Anatomically-constrained BIdirectional Connectivity” (GABIC).

The GABIC matrix is estimated from the NDTE flow, which is bi-directional, and is therefore asymmetric. This is unlike the anatomical matrix extracted by dMRI tractography which is un-directional. Importantly, GABIC is defined as the generators accounting for the causal influence of one neural system over another.

While there is a large literature on the dynamic causal modelling of effective connectivity within and among local neuronal (mass) models, the nonlinear and emergent dynamical properties of these systems have yet to be explored thoroughly. Previous models used the unidirectional structural connectivity (SC) to reproduce FC (Deco *et al.*, 2011; Deco and Kringelbach, 2014) and a common global conductivity value meaning that only the scaling factor is optimized. In other words, the scaling factor is a global coupling parameter expressing the conductivity of all fibers equally. In contrast, another possibility is to tune network connections individually (but not independently), which requires a dedicated estimation procedure (Gilson *et al.*, 2016). This corresponds to the concept of effective connectivity (EC) (Friston, 2011), which describes how network nodes excite or inhibit each other for a given model of local dynamics. EC thus describes causal interactions whose effects are modulated by the local dynamic regime of the node, which may shape FC in a complex fashion (Park and Friston, 2013): two areas may be significantly correlated (in FC) although disconnected (EC=0) when strong indirect pathways connect them (i.e., large network effect). Biologically, EC measures the strengths of connections, which depend not only on anatomical properties embodied in SC values (connection densities), but also on heterogeneities in synaptic receptors or neurotransmitters. Gilson and colleagues provided a solution for estimating EC from fMRI FC with information about the directed connectivity at the whole-brain level (divided in a parcellation of 70 brain regions with a couple of thousands connections to tune).

Here, we significantly extend existing models by using a more powerful bidirectional measure, namely the NDTE flow that properly captures the underlying spatiotemporal dynamical causal mechanisms, and thus not correlations or timeshifted correlations. For that we use a recent successful model, namely the Hopf whole-brain model (Deco *et al.*, 2017c), in combination with a powerful non-gradient based global optimization algorithm, namely the particle swarm optimization.

Briefly, in the following we describe how whole-brain models aim to balance between complexity and realism in order to describe the most important features of the brain *in vivo* (Cabral *et al.*, 2017). This balance is extremely difficult to achieve because of the astronomical number of neurons and the underspecified connectivity at the neural level. The emerging collective macroscopic dynamics of brain models use mesoscopic top-down approximations of brain complexity with dynamical networks of local brain area attractor networks (Breakspear, 2017; Deco and Jirsa, 2012). Essentially, these models link anatomical structure (given by the dMRI tractography) and functional dynamics (typically measured with fMRI) to reproduce the whole-brain empirical data (Deco *et al.*, 2015; Jirsa *et al.*, 2002).

Here we use the Hopf whole-brain model consisting of coupled dynamical units (ROIs or nodes) representing the N cortical and subcortical brain areas from a given parcellation (Deco *et al.*, 2017c). The local dynamics of each brain region is described by the normal form of a supercritical Hopf bifurcation, also known as the Landau-Stuart Oscillator, which is the canonical model for studying the transition from noisy to oscillatory dynamics (Kuznetsov, 1998). Coupled together with the brain network architecture, the complex interactions between Hopf oscillators have been shown to reproduce significant features of brain dynamics observed in electrophysiology (Freyer *et al.*, 2011; Freyer *et al.*, 2012), MEG (Deco *et al.*, 2017b) and fMRI (Deco *et al.*, 2017d; Kringelbach *et al.*, 2015).

The dynamics of an uncoupled brain region *n* is given by the following set of coupled dynamical equations, which describes the normal form of a supercritical Hopf bifurcation in Cartesian coordinates:

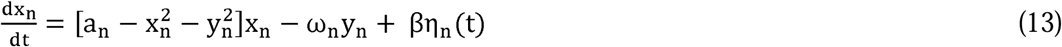

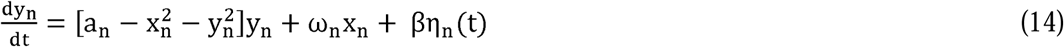

where η_n_(t) is additive Gaussian noise with standard deviation β. This normal form has a supercritical bifurcation a_n_ =0, so that if a_n_ >0, the system engages in a stable limit cycle with frequency *f*_*n*_ = ω_n_/2*π*. On the other hand, when a_n_<0, the local dynamics are in a stable fixed point representing a low activity noisy state. Within this model, the intrinsic frequency ω_n_ of each region is in the 0.008–0.08Hz band (*n*=1, …, *N*), where *N* is the total number regions.

We estimated the intrinsic frequencies from the empirical data, as given by the averaged peak frequency of the narrowband BOLD signals of each brain region. The variable x_n_ emulates the BOLD signal of each region *n*. To model the whole-brain dynamics we added an additive coupling term representing the input received in region *n* from every other region *p*, which is weighted by the corresponding structural connectivity. The whole-brain dynamics was defined by the following set of coupled equations:

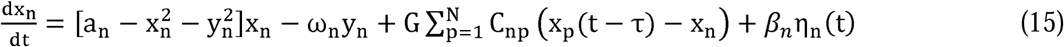

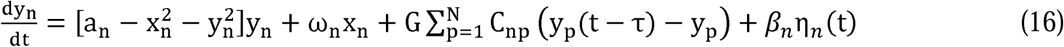

Where G denotes the global coupling weight, scaling equally the total input received in each brain area, and τ is a time lag. The initial values of noise was fixed thus: β=0.02, a_n_ = 0, and the time lags also initialized to zero. The structural connectivity matrix C_np_ is estimated and normalised from dMRI tractography (with a max of 0.2) and thus symmetric. We optimize sequentially, for each local region, the noise level, *β*_*n*_, the time lag, τ, the local bifurcation parameters, a_n_, and most importantly the matrix C_np_.

During optimisation, the strength of connections in C_np_ is updated based on the fit between the model output and the empirical NDTE flow matrix in terms of correlation. The empirical NDTE matrix is bidirectional and thus asymmetric. Hence, when updating the structural matrix, this will become asymmetric too. Thus C_np_ is the generative anatomically-constrained bidirectional connectivity, GABIC.

We used a global optimization routine of MATLAB, namely the particle swarm optimizer. Particle swarm is a population-based algorithm, similar to genetic algorithms (Kennedy and Eberhart, 1995). A population of individuals (called particles) diffuse throughout the searching region of parameters, not dissimilar to flocks of insects swarming. At each step, the algorithm evaluates the objective function for each particle. In our case, the objective function consisted of maximizing the correlation between the empirical and simulated NDTE matrix (i.e. considering the causality between all pairs). The diffusion of the particles is optimally adapted by the algorithm in order to converge to a global maximum.

## Acknowledgements

We would like to thank Daniel Chicharro, Jean-Pierre Changeux and Stanislas Dehaene for their insightful comments on the manuscript.

## Supplementary figures

**Figure S1.**
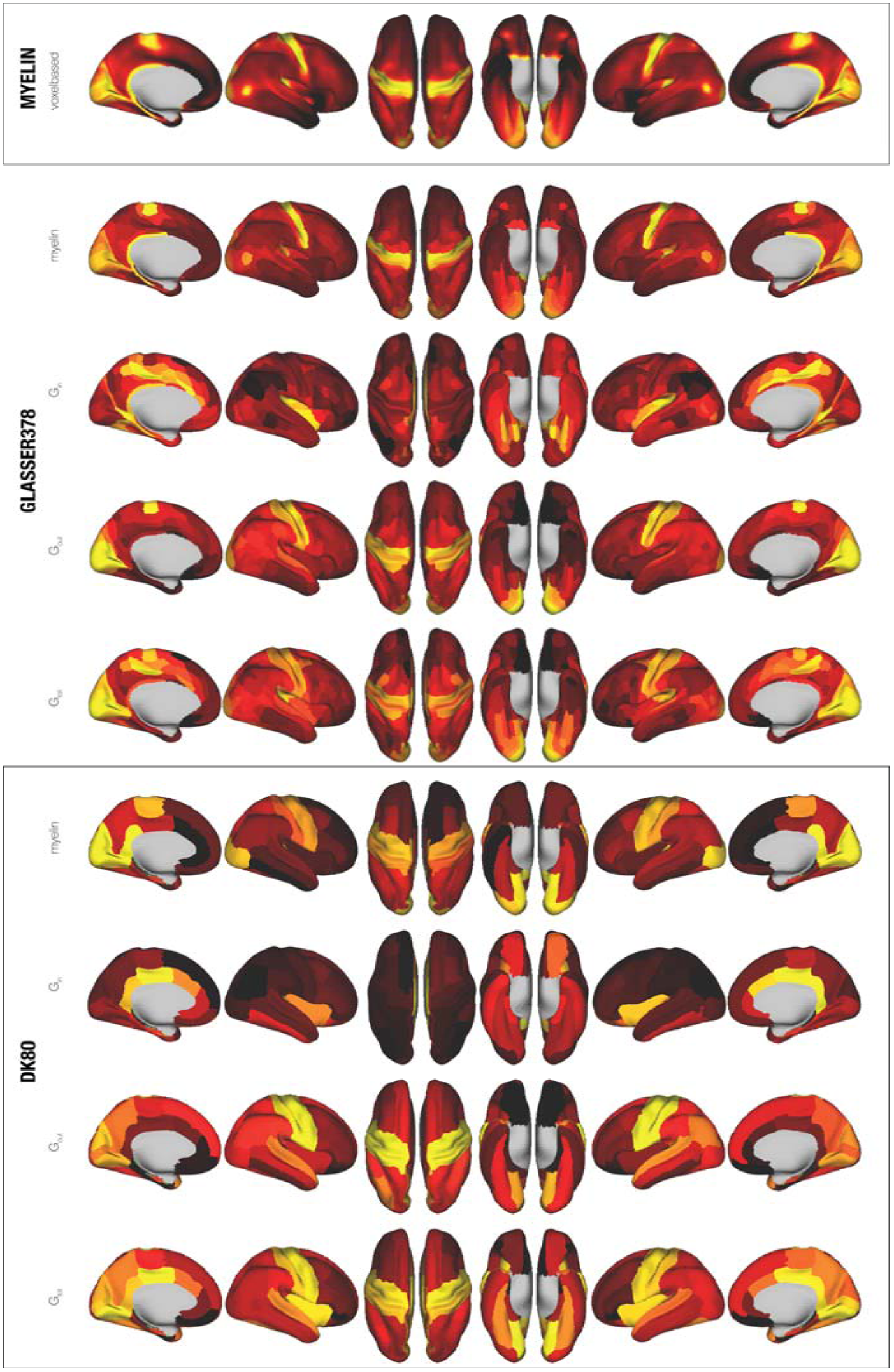
Functional hierarchy of resting state found using NDTE rendered on Glasser and DK80 parcellations. At the top is shown the full cortical voxelbased renderings of myelin (T1w/T2w). Below is shown renderings for both the Glasser and DK80 parcellations of myelin and the NDTE measures (G_in_, G_out_ and G_tot_) in 1003 HCP participants.

**Figure S2.**
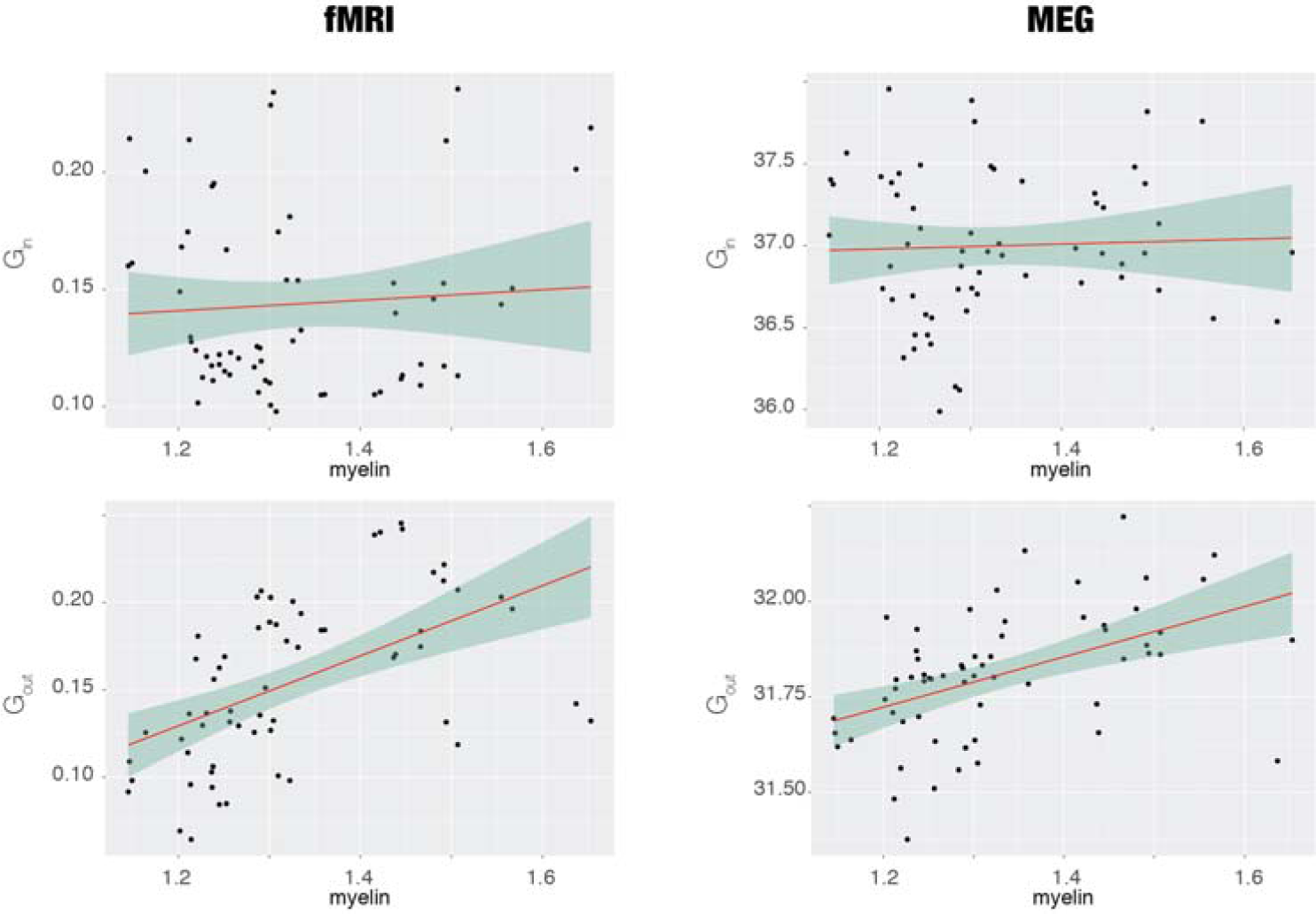
Validating NDTE framework using MEG from 89 HCP participants. We used the NDTE framework on MEG resting state data from 89 HCP participants (each having three resting state sessions) and extracted the timeseries from the 62 cortical regions of the DK80 parcellation (see Methods). On the left is shown the scatterplots for G_in_ and G_out_ for fMRI resting state data versus myelin (T1w/T2w). On the right is shown the same scatterplots for G_in_ and G_out_ for MEG resting state data versus myelin. The correlation values are 0.04 for G_in_ (14-22Hz, window size 1000ms), 0.48 for G_out_ (22.5-30.5Hz, window size 500ms).

**Figure S3:**
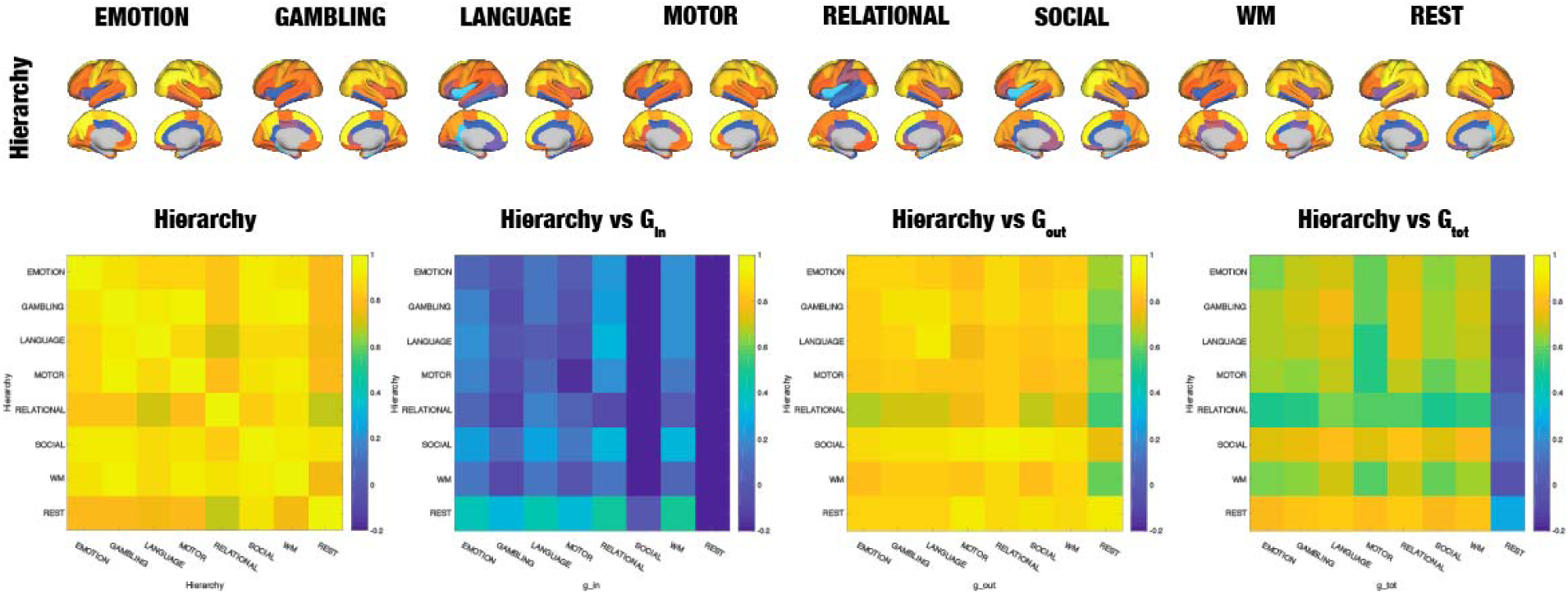
Alternative measure of hierarchy. Inspired by the neuroanatomical research by Markov and colleagues, we computed a similar measure of hierarchy as the fraction of feedforward and feedback (FF) organisation. We computed this for the seven tasks and for resting state (top row). As can be seen from the correlation matrices, this FF Hierarchy measure is not discriminatory between the seven tasks and only weakly correlated with, the integrative measure of incoming information flow.

